# Sublethal Behavioural and Neurotoxic Effects of Wastewater Effluent Exposure in a Freshwater Crustacean

**DOI:** 10.64898/2026.03.30.715298

**Authors:** Natalia Sandoval Herrera, Elin Johansson Kvarnström, Lea M. Lovin, Jerker Fick, Erin S. McCallum

## Abstract

The increasing discharge of treated wastewater effluent poses a growing threat to freshwater ecosystems. Although wastewater treatment plants reduce chemical pollution, they do not fully remove many biologically active compounds. Behavioural responses in aquatic organisms provide sensitive and ecologically relevant indicators of sublethal contaminant exposure, offering insight into underlying physiological disruption and potential ecological consequences. Here, we examined the behavioural and neurotoxic effects of a seven-day experimental exposure to treated wastewater effluent in the noble crayfish (*Astacus astacus*). We quantified four ecologically important behaviours: (1) shelter use, a key antipredator strategy, (2) food seeking, (3) the ability to detect and respond to wastewater-associated olfactory cues, and (4) locomotor activity was assessed across all behavioural contexts. Cholinesterase (ChE) activity was measured as a biomarker of neurotoxicity. Exposure to wastewater effluent significantly altered crayfish behaviour. Exposed individuals exhibited higher locomotor activity compared to controls, exposed crayfish avoided areas containing wastewater cues, spending less time near the effluent source. Similarly, ChE activity was significantly reduced in exposed crayfish, indicating neurotoxic effects. The concurrence of ChE inhibition and behavioural modification suggests that effluent-derived contaminants may interfere with neural signalling pathways underlying crayfish locomotion and habitat selection. Overall, our results demonstrate that short-term exposure to treated wastewater effluent can induce both neurochemical disruption and ecologically relevant behavioural changes in *A. astacus*. Such alterations may increase vulnerability to predation and influence population dynamics in effluent-receiving waters, highlighting the importance of integrating behavioural endpoints with mechanistic biomarkers in assessing sublethal impacts of wastewater contamination.

## 1. Introduction

Municipal wastewater treatment plants (WWTPs) are widespread anthropogenic stressor for aquatic environments (Hamdhani et al., 2020; Undeman et al., 2022). WWTPs can change the structure and flow of receiving environments, and they release effluents that can alter the thermal environment, reduce water clarity, and contain diverse forms of chemical pollution including nutrients, metals, pharmaceuticals, and microplastics (Hamdhani et al., 2020; Loos et al., 2013; Sun et al., 2019; Undeman et al., 2022). Although conventional wastewater treatment processes can be effective at reducing inputs of nutrients and certain pollutants to surface waters, they were not optimized to fully remove novel pollutants like pharmaceuticals and pesticides. Centralized wastewater treatment is also not globally used, with an estimated 48% of wastewater being released back to surface waters completely untreated (Jones et al., 2021). Widespread sampling campaigns have shown that surface waters near WWTPs contain multiple pollutants, and that aquatic organisms like fish and benthic macroinvertebrates bioconcentrate these compounds into their tissues (Cerveny et al., 2021; McCallum et al., 2017; Richmond et al., 2018). Moreover, in arid environments, under extreme heat events, and in artificially constructed receiving environments, wastewater effluents can be the dominant source of water flow, further concentrating the exposure that animals experience (Grabicova et al., 2017; Luthy et al., 2015; Zhi et al., 2020). As such, researchers and environmental managers have been concerned over the effects that wastewater effluents can have on aquatic wildlife living in habitats near WWTPs. Although wastewater effluent contains complex mixtures of biologically active contaminants capable of inducing neurotoxicity and affecting animal behaviour, most ecotoxicological assessments focus on chemical analysis and apical endpoints, with limited integration of ecologically relevant behavioural responses and mechanistic biomarkers (Bertram et al., 2022; Ford et al., 2021).

While acute toxicity has been widely studied, sublethal effects of wastewater exposure are increasingly recognized as ecologically important. Behaviour is a particularly sensitive endpoint, as it integrates underlying physiological disruption, such as altered endocrine and neural signalling, with higher-order ecological processes including predator–prey interactions, habitat selection, and reproductive success. (Amiard-Triquet, 2009; Ford et al., 2021). To date, select studies have evaluated the impacts of wastewater effluent on aquatic organism behaviour. In the laboratory, fish have shown dampened aggression, activity, and boldness after exposure to wastewater effluent (McCallum et al., 2017; Mehdi et al., 2022; Melvin et al., 2016). In aquatic invertebrates, damselfly larvae exposed to wastewater effluent had reduced activity but faster c-start escape responses (Späth et al., 2022). Likewise, exposed amphipods showed reduced activity and were faster to perform reproductive behaviours (Love et al., 2020). Less research has focused on how wastewater exposure can impact more complex behaviours like resource use and habitat choice. Exposure to wastewater and the pollutants within it may change the way animals interact with their environment by modulating how they detect or perceive environmental cues. For example, tadpoles exposed to wastewater effluent showed dampened responses to olfactory cues of predation, indicating impaired sensory performance in both studies (Heerema et al., 2018; Troyer and Turner, 2015). The pollutants in wastewater might also affect how animals react to environmental cues if exposure impacts their perception of threat or risk. For example, marbled crayfish exposed to a mixture of six psychoactive pharmaceuticals were more active and spent less time in a sheltered area compared to control crayfish (Hossain et al., 2021). Therefore, aquatic organisms in wastewater impacted environments may encounter altered sensory landscapes, which can affect habitat selection and resource utilization.

Despite growing recognition of behavioural sensitivity to pollution, few studies have integrated behavioural endpoints with mechanistic biomarkers of neurotoxicity in freshwater invertebrates exposed to treated wastewater effluent. Cholinesterase (ChE) activity is a widely used biomarker of neurotoxicity, as this enzyme plays a critical role in synaptic transmission and can be inhibited by various pollutants, including pesticide compounds like organophosphates and carbamates (Nunes, 2011; Vioque-Fernández et al., 2007). *In vitro* assays have shown that wastewater effluents directly inhibit acetylcholinesterase activity (Kienle et al., 2019; Neale et al., 2017a), and *Daphnia magna* exposed to hospital wastewater effluents showed reduced ChE activity (Afsa et al., 2022). However, the extent to which wastewater-induced changes in ChE activity correspond with ecologically relevant behavioural alterations in aquatic invertebrates remains poorly understood.

Here, we assessed how a one-week exposure to treated wastewater effluent affects behavioural responses of noble crayfish *Astucus astucus*, alongside how exposure impacted a common biomarker of neurotoxicity. Crayfish are benthic omnivores that function as keystone species and ecosystem engineers in many freshwater systems (Reynolds et al., 2013). Their survival depends heavily on shelter use as an antipredator strategy and on olfactory cues for habitat selection and resource detection, and they have been previously used as water quality sentinels in environmental monitoring (Antón et al., 2000; Martin, 2014; Reynolds et al., 2013). Specifically, we assessed 1) how wastewater exposure affected shelter-seeking and foraging responses; 2) if crayfish detected (and potentially preferred or avoided) olfactory cues of wastewater effluents; 3) how exposure affected crayfish locomotor activity across these behavioural contexts, and 4) how exposure affected the activity of cholinesterase, an enzyme and commonly used biomarker of neurotoxicity from pollutant exposure. We predicted that if crayfish could detect wastewater-associated cues, exposed individuals would show altered spatial preferences, potentially avoiding effluent-derived cues. Chemicals like pesticides and some pharmaceutical contaminants present in effluent can induce physiological stress, oxidative damage, and metabolic disturbances, we further predicted that wastewater exposure would reduce cholinesterase activity and locomotor behaviour. Finally, we expected that exposure would interfere with shelter-seeking and foraging behaviours either via direct effects on resource detection or risk perception, or indirectly via altered locomotion.

## 2. Methods

### 2.1 Study species and housing

For this study, 58 noble crayfish *(Astacus astacus)* were purchased from a commercial supplier (Bo Konsult Förvaltning AB, Heby, Sweden). Until the experimental phase, the crayfish were housed in a 1000 L ground water flow-through tank supplemented with aeration. Ground water supply in Umeå municipality is ∼8°C, with a pH range of ∼8.0 (Vakin 2025a). The crayfish were provided with clay tile pieces and PVC tubes as shelters such that there were more shelters available than crayfish in the tank to avoid aggression (Nyström et al., 2023). Crayfish were fed frozen green peas (Nyström et al., 2023) twice a week, and this feeding frequency was determined to be optimal because more frequent feeding led to overfeeding. The light-dark cycle was set at 12:12 hours, with a one-hour sunrise and sunset fade. All crayfish were individually numbered with stickers and transparent tape adhered to their backs (carapace) for later identification throughout the behavioural tasks. During tagging, the animals were sexed, inspecting differences in genital openings and appendages (i.e., ovopores and gonopores).

### 2.2 Wastewater effluent collection

Treated wastewater effluent was collected from the Umeå wastewater treatment plant (Öns reningsverk) on September 25, 2023. The plant uses conventional mechanical, chemical, and biological treatments to process wastewater from over 110,000 households before releasing the treated water into the Ume River (Vakin, 2025b). The wastewater effluent was collected in 25 L plastic containers and transported to the Swedish University of Agricultural Science in Umeå. Upon arrival, the effluent was refrigerated at 4°C and kept in dark conditions to prevent potential changes in chemical composition due to temperature or photodegradation.

### 2.3 Wastewater exposure

After tagging, the crayfish were split into two groups, aiming to have a similar number of males and females in each group, and then transferred them to one of four exposure tanks: two for wastewater effluent exposure (exposed to 100% treated wastewater) and two for control (filled with tap water). Group 1 had 14 control and 15 exposed individuals, and Group 2 had 15 control and 14 exposed individuals. As in the housing tank, each exposure tank was equipped with aeration and provided clay tile pieces and PVC tubes as shelters. The crayfish were exposed for seven days, with a two-day gap between each experimental group’s start to facilitate the logistics of running behavioural trials. On days 3 and 5, a 50% water or wastewater exchange was performed, and the crayfish were fed. See Figure 1 for an overview of the experimental timeline.

**Figure 1:**
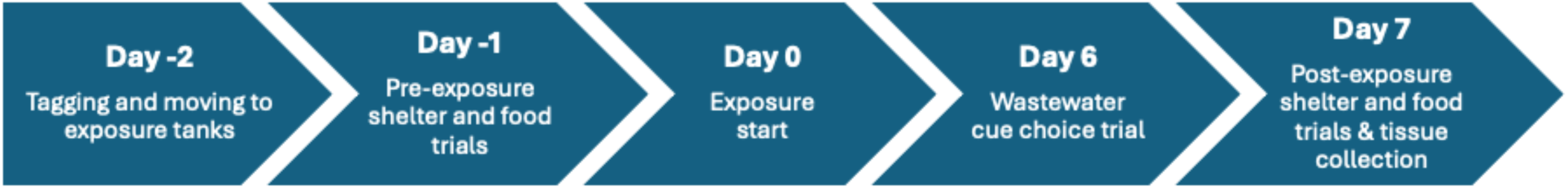
Experimental timeline showing the ordering of the behavioural tasks before and after seven days of wastewater effluent exposure. See Supplementary Figure S1A-C for schematics of behavioural trial tanks.

Water quality measurements were conducted on days 1, 3, 5, and 7. pH, conductivity (µS), total dissolved solids (TDS, ppm), and salinity (ppt) were measured with the Hach Pocket Pro+ Multi 2 Tester. Temperature and dissolved oxygen (DO, mg/L) with YSI Ecosense ODO200 Optical Dissolved Oxygen Meter. The eSHa Aqua quick test strip was used to measure total hardness (GH, mg/L CaCO_3_), carbonate hardness (KH, mg/L CaCO_3_), nitrite (NO_2_, mg/L), nitrate (NO_3_, mg/L), and chlorine (Cl_2_, mg/L). Water quality measures are summarised in Supplementary Table S1.

### 2.4 Behavioural trials

For the behavioural tasks, 50-L glass aquariums were used as experimental tanks. The aquarium walls were covered with adhesive white paper to prevent visual interaction among individuals. White aquarium gravel (∼1 cm) was added to the tank ground for traction and to ease contrast with the substrate for later behavioural scoring. The tasks were recorded from above using GoPro Hero 8 cameras.

### 2.4.1 Shelter-seeking trial

The shelter-seeking task was conducted twice: before and after exposure. In pre-exposure trials, aged tap water was used for both control and exposed individuals, while post-exposure trials used either wastewater effluent or aged tap water, depending on the exposure. A terracotta pot (11 cm diameter and 10 cm height) was provided in each experimental tank as a shelter, placed on the short side of the tank with the opening facing the centre (Fig. 2). The placement of the terracotta pot was switched between trials to prevent side biases. To begin a trial, crayfish were moved from the exposure tanks to holding tubes placed in the experimental tanks on the opposite side of the shelters. The holding tubes were made of white PVC with a diameter of 8 cm and had drilled holes for water exchange. The cameras were activated, and after 5 minutes of acclimation, the crayfish were released from their holding tubes to move freely for 5 minutes, which was recorded.

**Figure 2:**
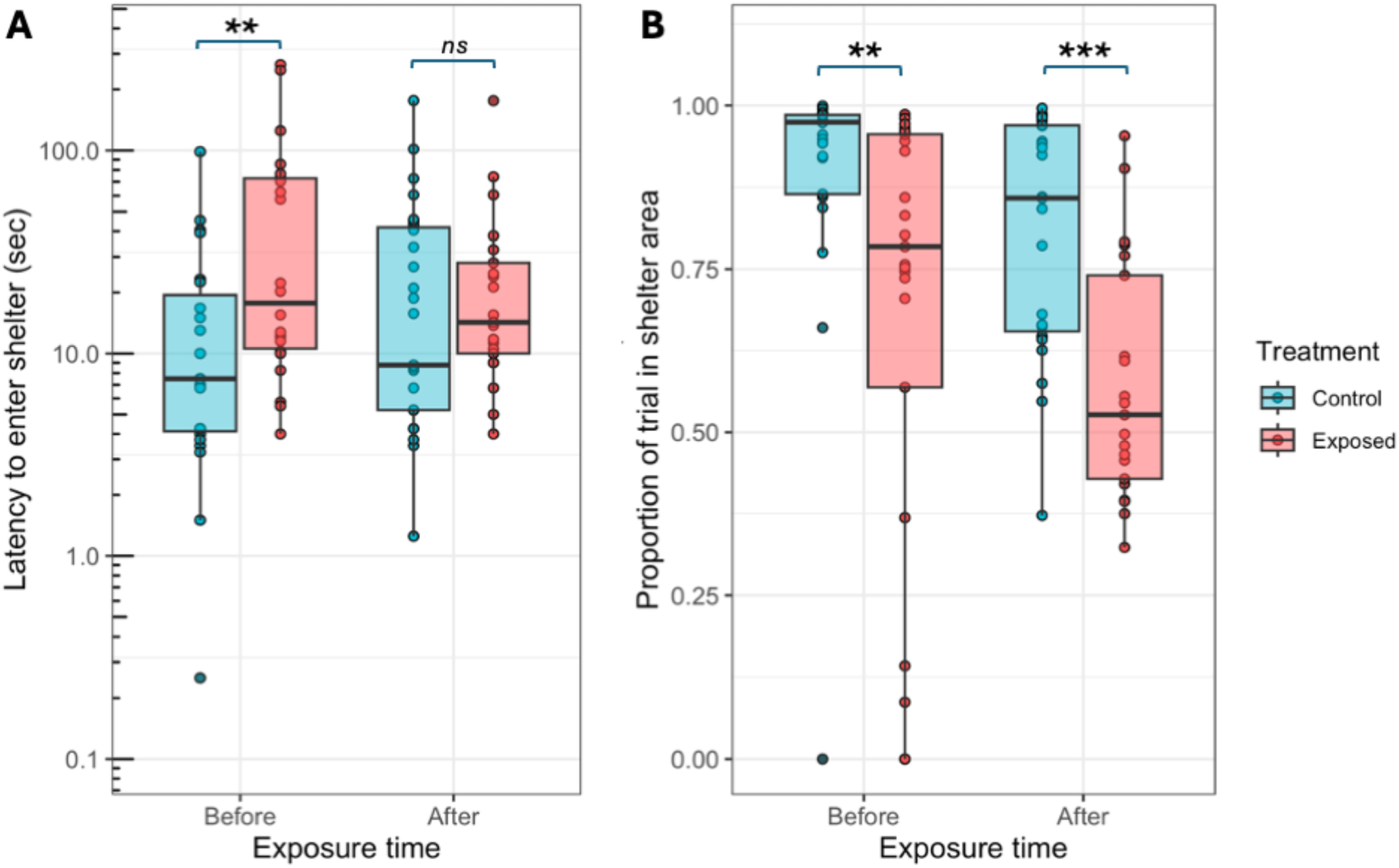
Effect of wastewater effluent exposure on crayfish shelter-seeking behaviour. A) Latency to enter the shelter area (seconds) plotted on a log_10_ axis. B) Proportion of trial time in the shelter area. Boxplots show the median and interquartile range overlaid on raw data points. Statistical significance stars show results from posthoc contrasts of treatment within exposure time. ** p < 0.01, *** p < 0.001, ns = not statistically significant, p > 0.05.

### 2.4.2 Food-seeking trial

The food-seeking task was conducted twice, like the shelter-seeking trial. In pre-exposure trials, aged tap water was used for both control and exposed individuals, while post-exposure trials used either wastewater effluent or aged tap water, depending on the treatment group. Five green peas were provided in each experimental tank as a food resource, placed on one side of the tank. Peas were used since this was the food during laboratory housing condition. The placement of the peas was counter-balanced across trials to prevent side biases. Trials were started and recorded for the same acclimation and duration as the shelter-seeking trials.

### 2.4.3 Wastewater cue choice trial

The wastewater cue preference task was performed at the end of the exposure period (day 6), before the final day when the shelter-seeking and food-seeking trials were repeated. The experimental tanks were filled with aged tap water, and PVC tubes of 1 cm diameter were attached to the short sides to deliver cues through peristaltic tubing. To begin a trial, crayfish were moved from the exposure tanks to holding tubes placed in the centre of the experimental tanks. The cameras were activated, and after 5 minutes of acclimation, the crayfish were released from their holding tubes to move freely for 15 minutes. The pumps delivering the cues were started after the crayfish were placed in their holding tubes. Each crayfish was presented with two cues, wastewater effluent or tap water, one on each side of the tank. Peristaltic pumps (Masterflex L/S, with Masterflex 06424-14 tubing) kept a constant flow rate of each cue at 6 mL/min which showed during a previous study and pilot trials with food colouring dye to have the cues meet after 15 min (Rossi, 2025). The delivery side of the wastewater cue was switched across trials to prevent side biases. The tanks were cleaned with tap water and 70% ethanol to eliminate residual cues between each set of trials.

### 2.5 Body size measurements and tissue sampling

Once the behavioural tasks were completed, the crayfish were euthanized with cold sedation followed by decapitation and photographed for later body size measures. Tissue sample (1 cm x 1 cm) was collected from the tail flexor muscles and stored frozen at -80°C until analysis. Total body length was measured from the top of the rostrum to the end of the telson using the Image J program (version 1.53m, 2024).

### 2.6 Cholinesterase neurotoxicity biomarker assay

Cholinesterase activity was measured in crayfish tail muscle using a colorimetric assay. Approximately 0.4 g (wet weight) of tissue was homogenized in a 1:10 (w:v) potassium phosphate buffer (0.1 mM, pH 7.2) for 30 s at 24,000 oscillations min⁻¹ with zirconium beads (Mini Beadbeater, BioSpec, Bartlesville, OK), then centrifuged at 1500 g for 5 min at 4 °C. The supernatant was used for the assay. First, protein concentration was determined by the Bradford assay (Sigma® reagent) with bovine serum albumin as the standard. Then, ChE activity was quantified by the Ellman method using acetylthiocholine (75 mM) as substrate and DTNB (10 mM) as chromogenic reagent. Absorbance was recorded at 412 nm for 15 min (BioTek Eon microplate spectrophotometer). Activity was expressed as U mg⁻¹ protein, where one unit corresponds to the hydrolysis of 1 µg of substrate per minute. All samples were run in triplicate and averaged to give the final value.

### 2.7 Chemical analysis of wastewater

We collected and froze water samples from both treatments across the study period to quantify a panel of 70 pharmaceuticals, personal care products, and pesticides in the effluent (*N* = 4 control treatment samples and *N* = 4 wastewater treatment samples, one from each experimental group at the start and end of the exposure). Water samples were frozen at -20°C until later analysis. A full list of analytes is given in Supplementary table S

We used offline solid phase extraction (SPE) followed by mass spectrometry to quantify concentrations of pollutants in the water samples following established protocols (Brand et al., 2024). Briefly, we thawed the water samples in the fridge overnight before extraction. Samples were extracted on an SPE manifold (Biotage VacMaster) using Oasis HLB cartridges (6 cc, 200 mg sorbent, 30 μm; Waters Corporation). The cartridges were pre-conditioned using 5 mL methanol (Merck, HPLC grade, CAS 67-56-1) followed by 5 mL Milli Q Ultra Pure Water. Twenty-five ng of a mixture of 17 isotopically labelled internal standards (Supplementary Table S3) was added to each sample and mixed before it was passed through the cartridge at a rate of ∼ 1 drop per second. The cartridges were eluted into 12 mL glass vials with 5 mL methanol (Merck, HPLC grade) followed by 5 mL ethyl acetate (Merck, HPLC grade, CAS 141-78-6), and the sample was then evaporated to dryness (Biotage Turbovap). Samples were then reconstituted in 150 μL of methanol (Merck, LCMS grade, CAS 67-56-1) acidified with 0.1% formic acid, transferred to 1.5 mL glass autosampler vials with a 200 μL glass insert, and frozen at -20°C until liquid chromatography mass spectrometry analysis.

Liquid chromatography mass spectrometry analysis was made by injecting 10 uL of each sample on a UHPLC system with an analytical column (Thermo Scientific Hypersil GOLD, 50 × 2.1 mm, 5 μm) equipped with a precolumn (Hypersil GOLD, 10 × 2.1 mm, 3 μm), connected to a TSQ Quantiva triple quadrupole mass spectrometer (Thermo Scientific) equipped with a heated-electrospray ionization ion source operating in positive mode. Chromatographic separation was achieved using a gradient of water, acetonitrile and methanol (LiChrosolv, Merck, Darmstadt, Germany). The resolution for both quadrupoles was 0.7 fwhm. Spray voltage was 3500 V, sheath gas 40 arbitrary units, sweep gas 0 arbitrary units, ion transfer tube temperature 350 °C, and vaporizer temperature 338 °C. For control of the mass spectrometer and data analysis Xcalibur (Thermo Scientific) was used. Two MS/MS transitions, one for quantification, one for qualification, were monitored for all analytes.

### 2.6 Video analysis of behaviour

The behaviour of crayfish during the experiments was analysed using the EthoVision XT software (version 16.0.1538, 2021). All videos were tracked at 4 frames per second. The program identified the crayfish in each video using automated dynamic background subtraction, and we further optimized crayfish identification with fine-scale contrast thresholding. The software tracked the crayfish to measure their distance moved and latency to reach or time spent in specific zones (e.g., inside shelters or sides of the wastewater cue zone, detailed below). All tracks were manually reviewed to identify program tracking errors (e.g., when the program failed to detect the crayfish). Errors were corrected manually with manual position assignment and track interpolation.

For shelter seeking trials, the shelter zone was a rectangle around the shelter clay pot that would capture when crayfish sought refuge in, behind, or under the edges of the pot (Supplementary Figure S2A). We measured latency to reach the shelter zone in seconds, proportion of the trial time spent in the shelter zone, and activity rate as distance travelled (cm per minute).

For food-seeking trials, the food resource zone was a circular zone drawn around the peas (Supplementary Figure S2B). We measured latency to reach the food zone in seconds and activity rate as distance travelled (cm per minute).

For the wastewater cue choice trial, the test tank was divided into six equal rectangular zones (Zones A-E). The wastewater cue preference zone was Zone A, closest to the wastewater cue inlet (Supplementary Figure S2C). We quantified preference as the proportion of time spent in Zone A on a per minute basis for each crayfish to capture how preference might change over time. We also measured activity rate as distance travelled over the entire trial (cm per minute).

### 2.7 Statistical analyses

All statistical analyses were conducted in R (version 4.5.0, 2025-04-11) (R Core Team, 2023). Model fit and assumptions using diagnostic plots and test provided in the packages performance (version 0.14.0) and DHARMa (version 0.4.7) (Hartig, 2019; Lüdecke et al., 2021). In all the below described statistical models, we evaluated if any two-way interactions among significantly improved model fit using drop1, and removed them if they did not or were not predicted *a priori* (described further below). All regression models were analyzed using the glmmTMB package (version 1.1.14) (Brooks et al., 2017) and posthoc contrasts were conducted using the emmeans package (version 1.11.1) (Lenth, 2023).

For the shelter-seeking behaviour trial, we analyzed two response variables in separate models. We analyzed latency to reach the shelter zone using a linear mixed effects model (LMM). We a log_10_-transformed the response variable to meet model assumptions. We included time (before, after), treatment (exposed, control), sex (male, female), and experimental group (one, two) as categorical predictors. We included a random intercept of crayfish ID to account for repeated measures from the same individual. Next, we analyzed the proportion of time spent in the shelter zone using a generalized linear mixed effects model (GLMM) assuming a beta distribution appropriate for a continuous proportion data bounded between 0 and 1. We included time, treatment, sex, and experimental group as categorical predictors. We included a random intercept of crayfish ID to account for repeated measures from the same individual. In both models, we retained the two-way interactions between treatment and time, as they were predicted *a priori*. We followed each model with posthoc contrasts to assess whether wastewater treatment affected the response variables within each time point (i.e., before, after).

For the food-seeking behavioural trial, we the probability that a crayfish would interact with the food during the trial as a binary response variable (yes, no). We included treatment, time, sex, and experimental group as categorical predictors. We included a random intercept of crayfish ID to account for repeated measures from the same individual. We retained the two-way interaction between treatment and time, and we used posthoc contrasts to assess whether wastewater treatment affected the response variable within each time point (i.e., before, after).

For the wastewater cue choice trial, we analyzed the proportion of time spent in the zone nearest to the wastewater outlet (see description above) over the trial time, that was divided in one minute time bins, using a GLMM assuming a beta distribution appropriate for a continuous proportion data bounded between 0 and 1. We included time bin as a continuous predictor, and treatment, sex, experimental group, and cue side (right, left) as categorical predictors. We retained a significant interaction between experimental group and cue side in the final model. We included a random slope of time bin and a random intercept of crayfish ID to account for multiple measures from the same individual.

To assess how exposure affected crayfish activity (cm moved per minute) across the three behavioural trials, we combined all activity data from the choice trial and the shelter-seeking and food-seeking behavioural trials after exposure. We used a LMM and included treatment, behavioural trial type (choice, shelter, food), sex, and experimental group as categorical predictors. We included total length as a continuous predictor to account for the fact that larger crayfish will travel further distances. We included a random intercept of crayfish ID to account for the individual being represented in multiple behavioural trials. We retained a significant interaction between treatment and experiment in the final model, and we used posthoc contrasts in emmeans to test if the treatment groups significantly differed within each behavioural experiment.

We assessed how exposure impacted cholinesterase (ChE) activity using a linear model with treatment, sex, and experimental group as categorical predictors. We then tested if cholinesterase activity predicted crayfish locomotor activity across the three behavioural trials using a LMM where activity rate (cm moved per minute) was the response variable, ChE activity was a continuous predictor, body size was a continuous covariate, and a random intercept of ID was included to account for multiple behaviour measures from each crayfish.

Finally, the mortality data were analysed using a binomial generalized linear model. The model included the effects of treatment, sex, and experimental group as categorical predictors, with mortality as a binary, yes-no response variable.

All model syntax and statistical output are provided in the supplementary materials (Supplementary Tables S3-S9). All sample sizes for each analysis are detailed in supplementary data file.

## 3. Results

### 3.1 Behavioural results

#### 3.1.1 Shelter seeking

Five crayfish did not enter the shelter area and were removed from analysis of latency to reach the shelter zone (three from the wastewater exposed group, two from the control group). Latency to reach the shelter showed a significant interaction between exposure time and treatment (Figure 2A, Linear mixed effects model (LMM), *N =* 93, (Est ± SE): -0.41 ± 0.20, *Z* = -2.10, *p* = 0.036). This was driven by the treatment groups differing in their latencies *before* the exposure was started (emmeans, before: control vs exposed, -0.52 ± 0.15, *t* = -3.41, *p* = 0.0010), and they did not differ after wastewater exposure (emmeans, after control vs exposed, -0.10 ± 0.15, *t* = -0.72, *p* = 0.48). Body length, sex, and experimental group did not affect the latency to enter the shelter zone (Supplementary table S4).

There was no interaction between exposure time and treatment in the proportion of trial time crayfish spent in the shelter zone (Figure 2B, GLMM beta, *N =* 98, (Est ± SE): -0.12 ± 0.38, *Z* = - 0.31, *p* = 0.76). In general, crayfish in the wastewater exposure group spent less time in the shelter both before and after exposure (emmeans, before: control vs. exposed, 0.92 ± 0.28, *Z* = 3.29, *p* = 0.001; after: control vs. exposed, 1.04 ± 0.27, *Z* = 3.88, *p* = 0.0001). Sex and experimental group did not affect the proportion of trial time spent in the shelter zone (Supplementary table S5).

#### 3.1.2 Food seeking

Crayfish did not reliably interact with the food stimulus during the trials, and we did not observe any crayfish eating the food stimulus. Of the 91 trials, crayfish entered the food zone during 55 trials and did not enter the food zone during 36 trials. There was no interaction between exposure time and treatment for the probability that a crayfish would interact with the food resource (Supplementary Figure S5; Binomial GLMM, *N =* 91, (Est ± SE): 1.55 ± 1.01, *Z* = 1.53, *p* = 0.13). Fewer crayfish in the wastewater exposure group interacted with the peas before exposure (posthoc: before - control vs. exposed, 1.66 ± 0.79, *Z* = 2.11, *p* = 0.035), but not after (after - control vs. exposed, 0.11 ± 0.73, *Z* = 0.16, *p* = 0.88). Sex and experimental group did not affect whether or not a crayfish would interact with the food resource (Supplementary Table S6).

#### 3.1.3 Wastewater cue preference

When presented with a cue of wastewater effluent or control water in a two-choice arena, wastewater exposed crayfish spent proportionately less of the total trial time near the source of the wastewater effluent compared to crayfish from the control treatment (Figure 3, GLMM beta family, *N*_obs_ = 688, *N*_crayfish_ = 47, (Est ± SE): -0.32 ± 0.13, *Z* = -2.42, *p* = 0.016). Preference for the wastewater zone did not change significantly with trial time (0.023 ± 0.016, *Z* = 1.43, *p* = 0.15) or crayfish sex (-0.20 ± 0.15, *Z* = -1.36, *p* = 0.17; Supplementary Table S7).

**Figure 3:**
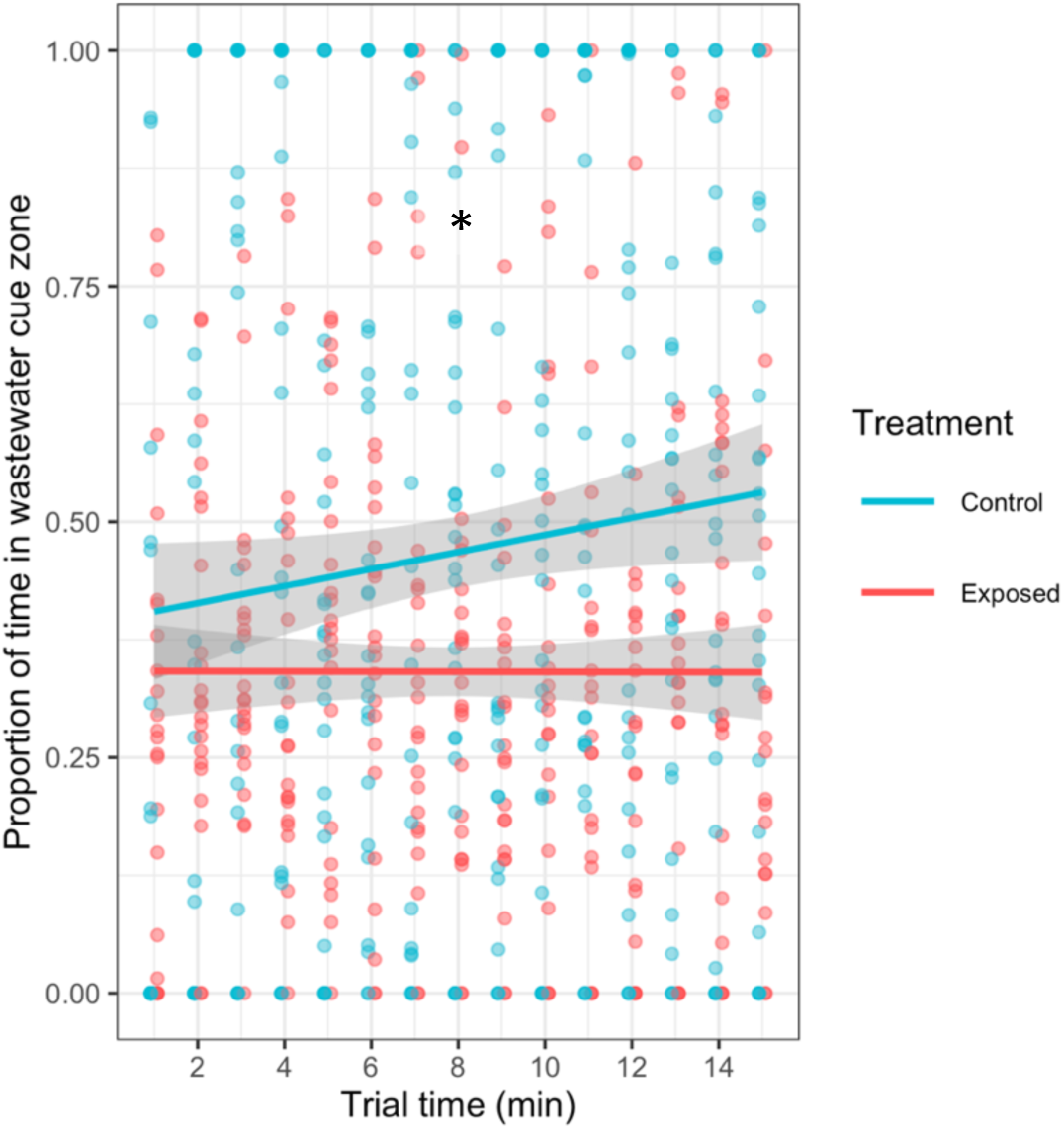
Effect of wastewater effluent exposure on crayfish preference for cues of wastewater effluent, shown as the proportion of time spent near the wastewater cue zone plotted by trial time (minutes). Solid line shows the average with the shaded ribbon showing the 95% confidence interval. *, p < 0.05.

#### 3.1.4 Activity rate across behavioural tasks

We measured activity across all three behavioural tasks and combined the data to assess how exposure to wastewater effluent affected crayfish activity rates across contexts. Wastewater treatment interacted with experiment, indicating that the effect of treatment depended on experiment. We compared treatments within experiment using posthoc contrasts to find that crayfish exposed to wastewater effluent were more active than those in the control treatment during the choice and shelter trials (Figure 4A, LMM, N_obs_ = 141; posthoc contrasts, Choice – control vs exposed (Est ± SE): -59.24 ± 8.39, *t* = -7.061, *p* < 0.0001; Shelter – control vs exposed: -22.83 ± 8.37, *t* = -2.73, *p* = 0.0073), but not for the food trial (Food – control vs exposed: -9.60 ± 8.56, *t* = -1.13, *p* = 0.26). Larger crayfish were more active (1.53 ± 0.56, *Z =* 2.71, *p* = 0.0067). Activity rates did not differ with sex or experimental group (all p > 0.05, Supplementary Table S8).

**Figure 4:**
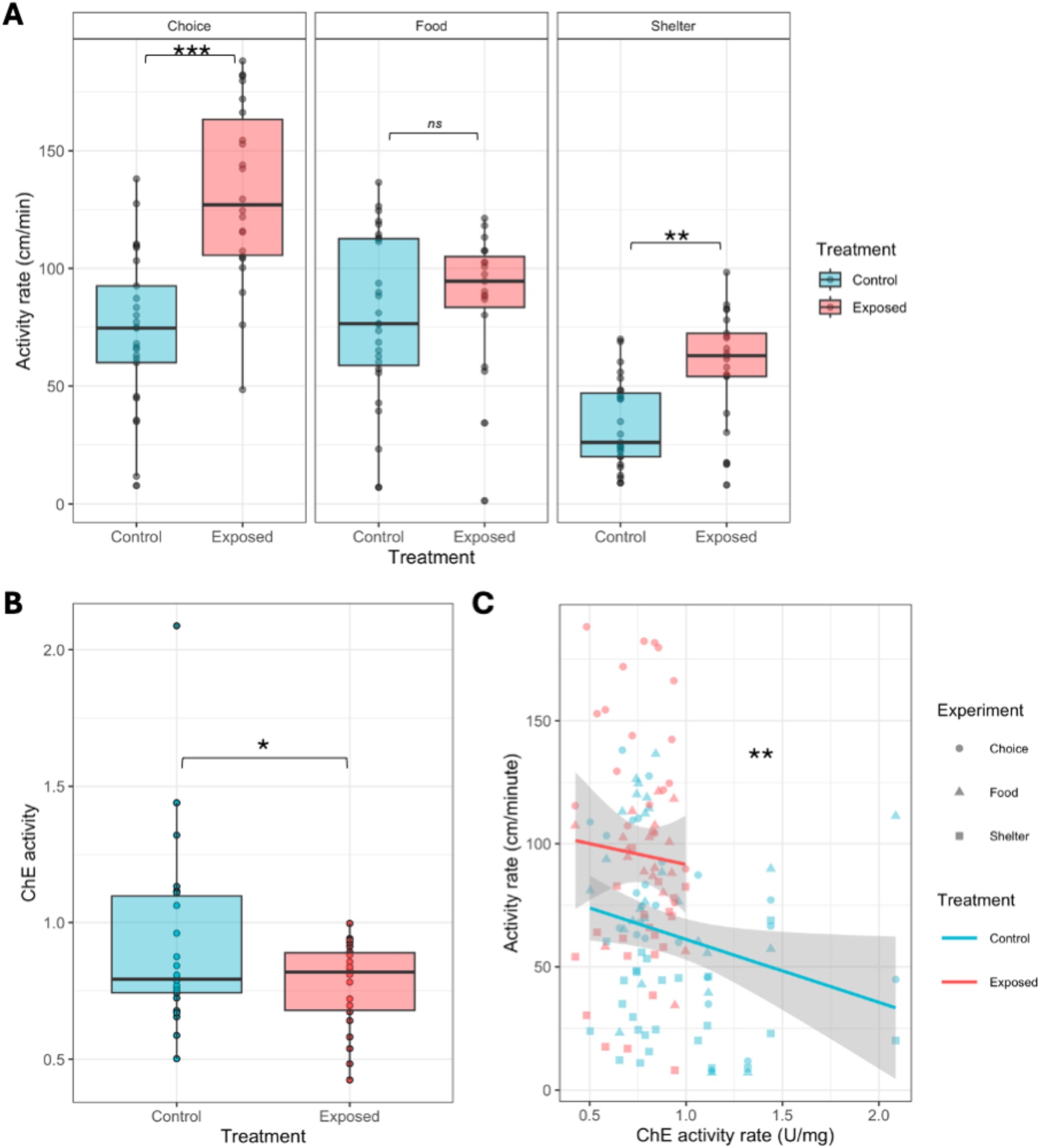
**A)** Effect of wastewater exposure treatment on locomotor activity rate (distance moved, cm per minute) faceted by the three behavioural trial types. **B)** Effect of wastewater treatment on cholinesterase (ChE) activity. **C)** Relationship between ChE activity and locomotor activity rate of crayfish during the choice task. For panels A and B, boxplots show the median and interquartile range, with raw data points plotted. For panel C, solid line shows the average value with shaded ribbons showing the 95% confidence interval, with raw data points plotted. * p < 0.05, ** p < 0.01, *** p < 0.001, *ns* = not statistically significant.

### 3.2 ChE neurotoxicity biomarker

Cholinesterase activity (ChE) was significantly lower in crayfish exposed to the wastewater treatment compared with unexposed controls (Figure 4B, Linear model, *N* = 48, (Est ± SE): -0.16 ± 0.078, *Z* = -2.07, *p* = 0.038). Exposed individuals had 17% lower ChE activity in their tail muscle, indicating potential neurotoxic impairment. There was no effect of sex or experimental group on ChE activity (Supplementary Table S9).

Cholinesterase activity significantly predicted locomotor activity rate in the choice across behavioural trials, where crayfish were more active with lower ChE activity (Figure 4C, LMM, *N = 137,* (Est ± SE): -36.51 ± 12.98, *Z* = -2.81, *p* = 0.0049; Supplementary Table S10)

### 3.3 Survival

We exposed 58 crayfish (39 males and 19 females). During the seven-day exposure period, nine crayfish died: seven from the exposed group (four males and three females) and two from the control group (both males). There was no significant effect of wastewater exposure treatment (Binomial generalised linear model [GLM], *N =* 58, (Est ± SE) 1.47 ± 0.85, *Z* = 1.73, *p* = 0.085); crayfish sex (GLM: -0.00023 ± 0.82, *z* = 0.00, *p* = 0.99) or experimental group (GLM: 0.33 ± 0.78, *Z* = 0.42, *p* = 0.67) on survival.

### 3.4. Water quality and chemical analysis of the wastewater effluent

Water quality measures are summarized in Supplementary table S1. The water in the wastewater exposure groups had higher salinity, conductivity, total dissolved solids, and carbonate hardness, while dissolved oxygen, temperature, pH, chlorine, nitrate, nitrite, and general hardness did not vary between the treatment groups.

We measured the concentrations of 70 pharmaceuticals, personal care products, and pesticides in the wastewater effluent (Figure 5). The compound with the highest concentration was the artificial sweetener acesulfame.

**Figure 5:**
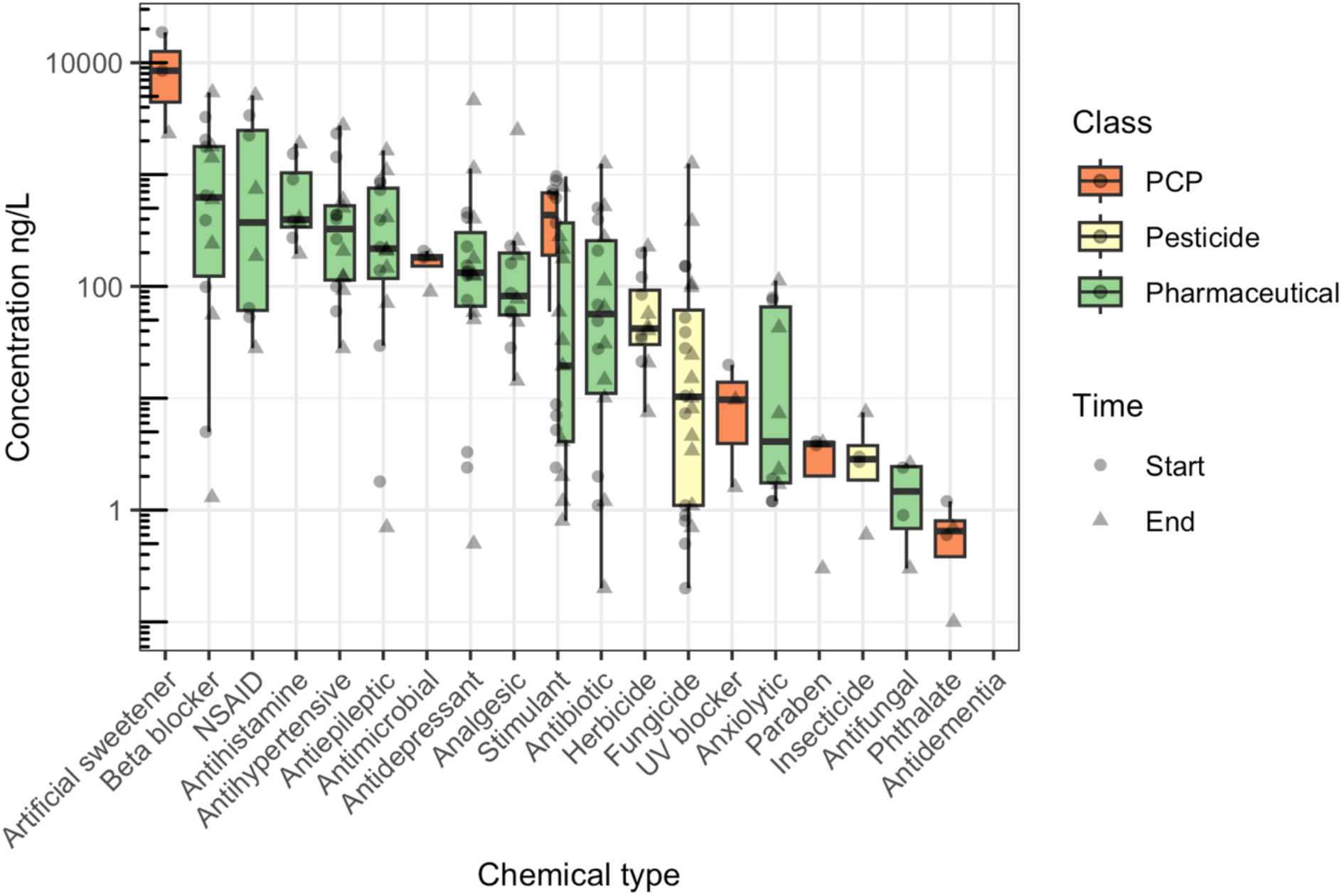
Median concentrations of chemical compounds measured in water samples collected from the wastewater exposed group at the start (*N* = 2, circular data points) and end (*N* = 2, triangular data points) of the exposure plotted on a log_10_ axis. Chemicals are grouped by chemical type (i.e., there are multiple chemicals within each type) and boxplots are coloured by compound class. See Supplementary table S2 for a full list of chemicals their groupings.

## 4. Discussion

We investigated how a one-week exposure to treated wastewater effluent affects behavioural responses of noble crayfish *Astucus astucus*, alongside how exposure impacted cholinesterase activity, a common biomarker of neurotoxicity. Specifically, we focused on how exposure affected shelter seeking, foraging, attraction to olfactory cues of wastewater, and locomotor activity across these contexts. We found that exposure to treated wastewater effluent increased crayfish activity across the three behavioural contexts, most notably in the shelter-seeking and cue preference trials. Prior research has shown that exposure to wastewater effluent has decreased locomotor activity in freshwater aquatic invertebrates or showed no effect on activity. For instance, *Gammarus pulex* exposed to 100% wastewater effluent decreased in activity over three weeks of exposure (Love et al., 2020), and damselfly larvae exposed to 100% treated wastewater effluent for one week moved slower and made more failed foraging attempts (Späth et al., 2022). However no effect has also been reported: *Gammarus fossarum* showed no change in locomotor activity after exposure to 100% effluent for 22 days (Rothe et al., 2022), and isopods (*Assellus a aquaticus*) exposed effluents showed no different in the percent of trial spent immobile after exposure for 21 days (Plahuta et al., 2017). We found no previous studies on the impacts of municipal wastewater effluents on crayfish behaviour, but several prior studies found that pharmaceuticals commonly measured in wastewater effluents can modulate crayfish activity. For instance, the anxiolytic oxazepam (∼700 ng/L) and the antidepressant sertraline (∼800 - 1000 ng/L) increased activity in two studies on the marbled crayfish (*Procambarus virginalis*) (Hossain et al., 2019; Kubec et al., 2019) and the swamp crayfish (*Procambarus clarkii*) (Suryanto et al., 2023), while the opioid analgesic tramadol (∼1 μg/L) reduced locomotor activity in marbled crayfish (Buřič et al., 2018).

The increased locomotor activity observed in exposed crayfish may reflect a stress-induced or neurophysiologically mediated response. Hyperactivity has frequently been reported following exposure to neuroactive contaminants, particularly those targeting cholinergic signalling pathways (Fulton and Key, 2001; Jebali et al., 2013). ChE is a well-established biomarker of neurotoxicity in aquatic organisms, particularly in response to organophosphate and carbamate contaminants (Nunes, 2011; Santana et al., 2021). However, more recently, some studies have shown that other compounds such as metals (Diamantino et al., 2003; Jebali et al., 2006), surfactants (Guilhermino et al., 2000), hydrocarbons (Oropesa et al., 2007), and pharmaceuticals (Nunes, 2011) could be sources of disruption of ChE function, most often at high contamination levels. As such, recent studies have shown that wastewater effluent exposure can inhibit ChE activity (Afsa et al., 2022; Neale et al., 2017b). Inhibition of this enzyme impairs the breakdown of the neurotransmitter acetylcholine, leading to its accumulation within synaptic clefts and subsequent overstimulation of cholinergic receptors. This overstimulation can initially manifest as increased neural excitation and erratic or elevated locomotor activity, followed by impaired coordination or fatigue at higher levels of inhibition. Previous studies in aquatic organisms further support the mechanistic linkage between ChE inhibition and altered behavioural performance. For instance, reductions in ChE activity have been directly associated with changes in swimming behaviour and escape performance in fish (Almeida et al., 2010; Rosales-Pérez et al., 2022; Sandoval-Herrera et al., 2019)(Santana et al., 2021), molluscs (Cooper and Bidwell, 2006), and crustaceans (reviewed in (Xuereb et al., 2009), indicating that disruption of cholinergic signaling can translate into ecologically relevant impairments in movement. Together, these findings suggest that contaminants found in the effluent here, many of which include pharmaceuticals, pesticides, and other neuroactive substances, may interfere directly with neurotransmission pathways, at environmentally relevant concentrations.

Crayfish that were exposed to wastewater effluent were more likely to avoid an olfactory cue of wastewater after exposure, spending ∼30% of the trial in the wastewater cue zone while control crayfish spent more time in this zone. Crayfish are highly sensitive to chemical olfactory cues that are sensed with antennules, located under their antennae (Derby, 2021). As an omnivore scavenger, detecting organic chemical cues is used in foraging and to detect predators (Derby, 2021; Reynolds et al., 2013). Because of this sensitivity, we hypothesized that crayfish would be able to detect wastewater effluent but were uncertain if they would be attracted to or avoid the stimulus. For instance, prior work found that marbled crayfish were attracted to cues of the antidepressant sertraline at ∼400 ng/L, and the authors concluded that sertraline might mimic natural chemical cues signaling food (Woodman et al., 2016). However, wastewater is a complex mixture of many compounds that may trigger an avoidance response, especially for particularly high chemical concentrations. For example, the amphipod *Gammarus fossarum,* behaviourally avoided spikes of copper and the insecticide methomyl and actively avoided wastewater effluents containing these compounds (Ruck et al., 2023). Our results suggest that crayfish can detect cues of wastewater effluent but may need prior exposure to form an association between olfactory cues of wastewater before initiating an avoidance response.

Wastewater exposure did not impact shelter-seeking or foraging behaviours. Exposed crayfish spent less time in the shelter both before and after exposure. Regardless of treatment, crayfish were quick to enter the shelter in the novel behavioural trial arena and often spent more than half of the trial time in the shelter, indicating that wastewater exposure may not affect their ability to identify the shelter as a refuge or change their perception of environmental threat such that they were less likely to use the shelter (e.g., if exposed to antidepressant or anti-anxiety pharmaceuticals). Unlike the shelter resource, crayfish did not reliably interact with or consume the food resource during the food seeking trial and wastewater exposure did not affect whether the crayfish would interact with the food. The crayfish were familiar with the food resource from its use prior to and during the exposure. We therefore hypothesize that crayfish may have needed a longer trial time to initiate a foraging response or they simply were not hungry, as they were fed during the exposure period.

In summary, we tested how a one-week exposure to treated wastewater effluent affects behavioural responses of freshwater noble crayfish *Astucus astucus*, alongside how exposure impacted a cholinesterase activity as common biomarker of neurotoxicity. The hyperactivity behavioural response correlated with a decrease in cholinesterase activity suggests that wastewater effluent, and the chemicals within it, inhibited cholinesterase activity at an intermediate level. The behavioural avoidance of wastewater cues by crayfish suggests that wastewater inputs may shape habitat choice and distribution of crayfish in the wild. Future research with longer exposure times would reveal how prolonged depression of cholinesterase and hyperactivity affects crayfish body condition, reproduction, and survival. Likewise, investigating how crayfish select and disperse from habitats with and without wastewater inputs would inform how pollution inputs like wastewater may shape aquatic invertebrate habitat choice and community composition in the wild.

## Credit author statement

**Natalia Sandoval Herrera**: Conceptualization, Formal analysis Investigation, Supervision, Writing – Review & Editing. **Elin Johansson Kvarnström**: Investigation, Formal analysis, Visualization, Writing – Review & Editing. **Lea M. Lovin**: Investigation, Writing – Review & Editing. **Jerker Fick**: Methodology, Resources, Writing – Review & Editing. **Erin S. McCallum**: Conceptualization, Formal analysis, Investigation, Resources, Writing – Original Draft, Visualization, Supervision, Project administration, Funding acquisition.

## Funding statement

This research was funded by the Swedish Research Council Formas grant (number 2020-00981) to E. McCallum and a Carl Tryggers Foundation grant (number 22:2082) to E. McCallum and L. Lovin.

## Data availability statement

All data associated with this project will be posted on the Open Science Framework (OSF) repository upon publication.

## Notes

### Competing Interest Statement

The authors have declared no competing interest.

