## Supplementary materials for "Sublethal Behavioural and Neurotoxic Effects of Wastewater Effluent Exposure in a Freshwater Crustacean"

### Supplementary Figures

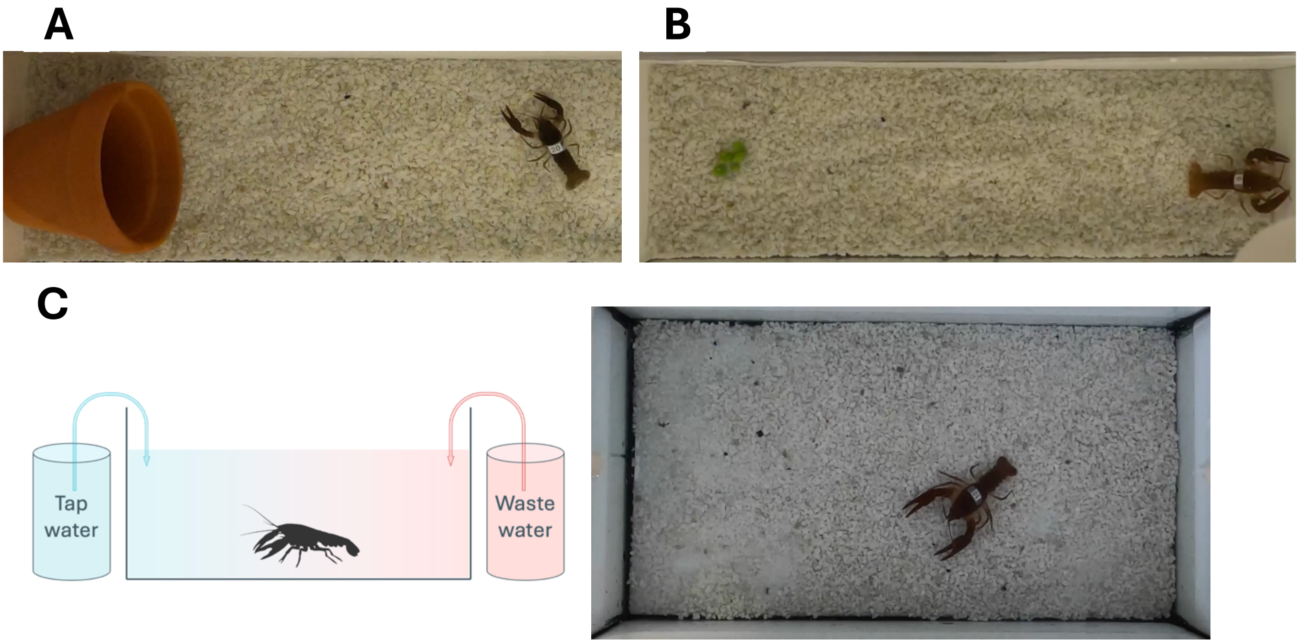


**Supplementary Figure S1**: Photographs of behavioural testing tanks for **A)** the shelter-seeking trial, **B)** the foraging trial, and **C)** the wastewater cue preference trial, including schematic showing cue delivery tanks.

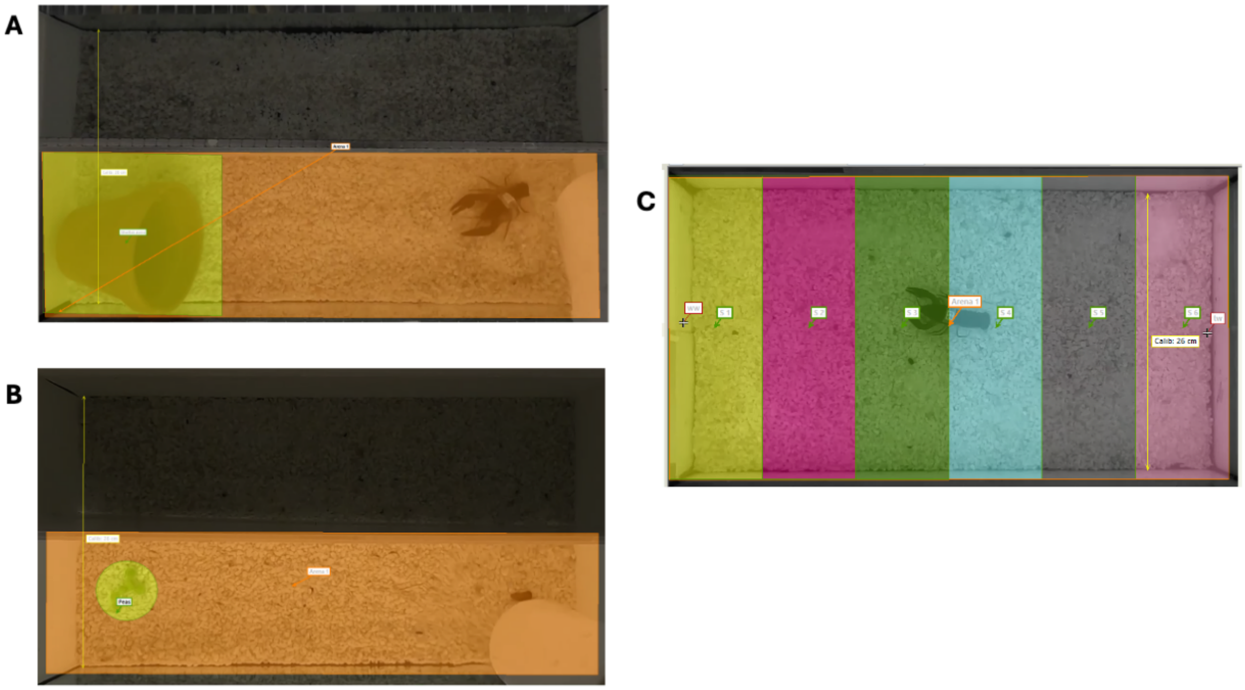


**Supplementary Figure S2**: Ethovision behaviour tracking arenas with zones for the three behavioural trials. A) The shelter seeking trial, with the shelter zone shown in yellow. B) The foraging trial, with the food zone shown in yellow. C) the wastewater cue preference trial, with the arena divided into six equal zones where zone S1 is closest to the wastewater cue. In all panels, a calibration bar is shown in yellow.

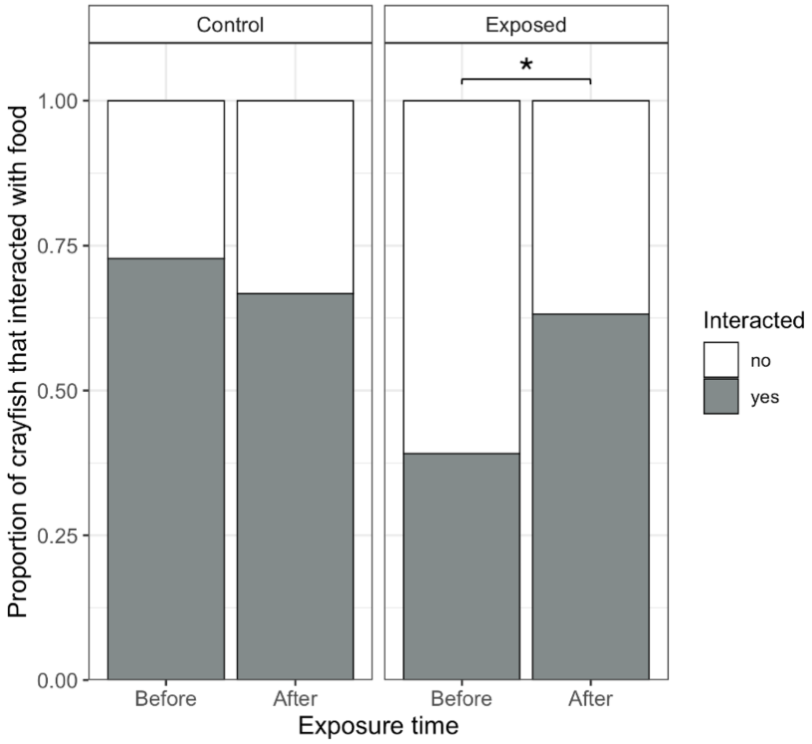


**Supplementary Figure S3:** Proportion of crayfish that interacted with the food stimulus during the foraging behavioural trial. Exposed crayfish interacted with the food more after exposure, but this did not differ from controls. * $p < 0.05$.

### Supplementary Tables

**Supplementary Table S1:** Water quality measured taken in exposure tanks across the study (mean ± standard deviation). Dissolved oxygen and temperature were measured on a YSI Ecosense. pH, salinity, conductivity, and total dissolved solids (TDS) were measured with a Hach multimeter. We measured carbonate hardness using eSHa 6-in-1 water test strips. This test also monitored chlorine (0 ± 0), nitrate (0 ± 0), nitrite (0 ± 0), and general hardness (~6 dH ± 0), but we never measured variation among the tests. Measures were taken on day 1, 3, 5 for experimental group 1, and 1, 3, 5, and 7 for experimental group 2.

| | DO mg/L | Temp °C | pH | Salintiy ppt | Conductivity µS | TDS ppm | KH °dH |
|---|---|---|---|---|---|---|---|
| Control *N* = 7 | 9.99 ± 0.64 | 16.63 ± 0.85 | 7.97 ± 0.13 | 0.081 ± 0.035 | 168.80 ± 72.35 | 119.81 ± 51.14 | 5.14 ± 0.90 |
| Exposed *N* = 7 | 9.41 ± 0.69 | 16.33 ± 0.76 | 7.80 ± 0.36 | 0.44 ± 0.029 | 885.86 ± 61.97 | 629 ± 43.93 | 13.43 ± 1.62 |

**Supplementary table S2**: Chemicals analytes, their classification by class and type of chemical, and their limit of quantification (LOQ). PCP = personal care product.

| Analyte | Class | Class specific | LOQ, ng/L |
|---|---|---|---|
| Acesulfame | PCP | Artificial sweetener | 5 |
| Amitriptyline | Pharmaceutical | Antidepressant | 5 |
| Amoxicillin | Pharmaceutical | Antibiotic | 5 |
| Amphetamine | Pharmaceutical | Stimulant | 0.1 |
| Atenolol | Pharmaceutical | Beta blocker | 1 |
| Atrazine | Pesticide | Herbicide | 5 |
| Atrazindesethyl | Pesticide | Herbicide | 5 |
| Azithromycin | Pharmaceutical | Antibiotic | 0.1 |
| Azoxystrobin | Pesticide | Fungicide | 0.5 |
| BAM | Pesticide | Herbicide | 5 |
| bentazon | Pesticide | Herbicide | 0.1 |
| Bisoprolol | Pharmaceutical | Beta blocker | 0.5 |
| Bixafen | Pesticide | Herbicide | 5 |
| Benzoylecognine | Pharmaceutical | Stimulant | 0.1 |
| Boskalid | Pesticide | Fungicide | 10 |
| Caffeine | PCP | Stimulant | 1 |
| Carbamazepine | Pharmaceutical | Antiepileptic | 0.1 |
| Cetirizine | Pharmaceutical | Antihistamine | 1 |
| Ciprofloxacin | Pharmaceutical | Antibiotic | 1 |
| Citalopram | Pharmaceutical | Antidepressant | 0.1 |
| Clarithromycin | Pharmaceutical | Antibiotic | 0.1 |
| Clindamycin | Pharmaceutical | Antibiotic | 0.1 |
| Clotrimazole | Pharmaceutical | Antifungal | 0.1 |
| Cocaine | Pharmaceutical | Stimulant | 0.5 |
| Codeine | Pharmaceutical | Analgesic | 1 |
| Cotinine | Pharmaceutical | Stimulant | 1 |
| DEHP; Bis(2-ethylhexyl) phthalate | PCP | Phthalate | 0.1 |
| Diazepam | Pharmaceutical | Anxiolytic | 1 |
| Diclofenac | Pharmaceutical | NSAID | 0.5 |
| erythromycin | Pharmaceutical | Antibiotic | 1 |
| Fexofenadine | Pharmaceutical | Antihistamine | 0.5 |
| Fluoxetine | Pharmaceutical | Antidepressant | 1 |
| Fluopyram | Pesticide | Fungicide | 0.1 |
| Gabapentin | Pharmaceutical | Antiepileptic | 0.1 |
| Ibuprofen | Pharmaceutical | NSAID | 1 |
| Imidacloprid | Pesticide | Insecticide | 1 |
| Irbesartan | Pharmaceutical | Antihypertensive | 0.5 |
| Isoproturon | Pesticide | Herbicide | 5 |
| Lamotrigine | Pharmaceutical | Antiepileptic | 0.5 |

| | | | |
|---|---|---|---|
| Lidocane | Pharmaceutical | Analgesic | 5 |
| Losartan | Pharmaceutical | Antihypertensive | 1 |
| Mekoprop | Pesticide | Herbicide | 5 |
| Memantin | Pharmaceutical | Antidementia | 0.1 |
| Metalaxyl | Pesticide | Fungicide | 0.1 |
| Methamphetamine | Pharmaceutical | Stimulant | 0.1 |
| Methylparaben | PCP | Paraben | 0.1 |
| Methylphenidate | Pharmaceutical | Stimulant | 0.1 |
| Metoprplol | Pharmaceutical | Beta blocker | 0.5 |
| Metribuzin | Pesticide | Herbicide | 1 |
| Mirtazapine | Pharmaceutical | Antidepressant | 0.1 |
| Octocrylene | PCP | UV blocker | 0.1 |
| Ofloxacin | Pharmaceutical | Antibiotic | 1 |
| Oxazepam | Pharmaceutical | Anxiolytic | 0.1 |
| Paracetamol | Pharmaceutical | Analgesic | 1 |
| Pirimicarb | Pesticide | Insecticide | 0.1 |
| Pregabalin | Pharmaceutical | Antiepileptic | 0.1 |
| Propioconazole | Pesticide | Fungicide | 0.1 |
| Protioconazole-destio | Pesticide | Fungicide | 0.1 |
| Quinmerac | Pesticide | Herbicide | 0.1 |
| Sotalol | Pharmaceutical | Beta blocker | 0.1 |
| Sulfamethoxazole | Pharmaceutical | Antibiotic | 0.5 |
| Tebuconazole | Pesticide | Fungicide | 0.1 |
| Telmisartan | Pharmaceutical | Antihypertensive | 0.1 |
| Temazepam | Pharmaceutical | Anxiolytic | 0.1 |
| Thiacloprid | Pesticide | Insecticide | 5 |
| Tramadol | Pharmaceutical | Analgesic | 0.1 |
| Triclosan | PCP | Antimicrobial | 1 |
| Trimethoprim | Pharmaceutical | Antibiotic | 0.1 |
| Valsartan | Pharmaceutical | Antihypertensive | 1 |
| Venlafaxine | Pharmaceutical | Antidepressant | 0.1 |

**Supplementary Table S3**: List and CAS IDs of 17 internal standards used in the measurement of chemicals in the water samples.

| Internal standard | CAS ID |
| --- | --- |
| Caffeine 13C3 | 78072-66-9 |
| Carbamazepine 13CD6 | 132183-78-9 |
| Carbaryl D3 | 1433961-56-8 |
| Carbofuran D3 | 1007459-98-4 |
| Codeine D6 | 1007844-34-9 |
| Dichlorvos D6 | 203645-53-8 |
| Fluconalzole D4 | 1124197-58-5 |
| Ibuprofen D3 | 121662-14-4 |
| Ketoprofen D3 | 159490-55-8 |
| Methylparaben D4 | 362049-51-2 |
| Metoprolol D7 | 1219798-61-4 |
| Ofloxacin D3 | 1173147-91-5 |
| Parathion D10 | 350820-04-1 |
| Propoxur D7 | 2140327-65-5 |
| Risperidone D4 | 1020719-76-9 |
| Tramadol 13C D3 | 1884682-17-0 |
| Trimethoprim D9 | 1189460-62-5 |

**Supplementary table S4:** Statistical model syntax and output for analysis of latency to enter the shelter zone during the shelter seeking behavioural trial.

| glmmTMB(log10(Latency to shelter zone) ~ (Time * Treatment) + Sex + Experimental group + Total length + (1\|ID), family = "gaussian", data = joined_sheldata) | | | | |
|---|---|---|---|---|
| | Estimate | Std. Error | *Z* | *P* |
| Time | 0.22 | 0.13 | 1.68 | 0.092 |
| Treatment (exposed) | 0.52 | 0.15 | 3.41 | **0.00066** |
| Sex (male) | -0.14 | 0.13 | -1.08 | 0.28 |
| Exposure group (two) | -0.049 | 0.12 | -0.41 | 0.68 |
| Total length | 0.0071 | 0.0098 | 0.72 | 0.47 |
| Time * Treatment | -0.41 | 0.20 | -2.10 | **0.036** |
| posthoc contrasts: emmeans (model, pairwise ~ Treatment \| Time) | | | | |
| | Estimate | Std. Error | *t* | P |
| Before: Control vs Exposed | -0.52 | 0.15 | -3.41 | **0.0010** |
| After: Control vs Exposed | -0.11 | 0.15 | -0.72 | 0.48 |

**Supplementary table S5**: Statistical model syntax and output for analysis of the proportion of trial time in the shelter zone during the shelter seeking behavioural trial.

$N_{obs}$ = 98, $N_{crayfish}$ = 56

| glmmTMB(Proportion time in shelter zone ~ (Time * Treatment) + Sex + Group + (1\|ID), family = "ordbeta", data = joined_sheldata) | | | | |
|---|---|---|---|---|
| | Estimate | St. Error | Z | P |
| Time | -0.59 | 0.27 | -2.14 | **0.032** |
| Treatment (exposed) | -0.92 | 0.28 | -3.24 | **0.0012** |
| Sex (male) | 0.079 | 0.21 | 0.38 | 0.71 |
| Exposure group (two) | 0.30 | 0.20 | 1.52 | 0.13 |
| Time * Treatment | -0.14 | 0.38 | -0.38 | 0.71 |
| posthoc contrasts: emmeans(model, pairwise ~ Treatment \| Condition) | | | | |
| | Estimate | St. Error | *Z* | P |
| Before: Control vs Exposed | 0.92 | 0.28 | 3.25 | **0.0012** |
| After: Control vs Exposed | 1.06 | 0.27 | 3.97 | **0.0001** |

**Supplementary table S6**: Statistical model syntax and output for analysis of the number of crayfish that interacted with the food resource during the food seeking behavioural trial.

$N_{obs}$ = 91, $N_{crayfish}$ = 56

| glmmTMB(Interacted with food YN ~ (Time * Treatment) + Sex + Group + (1\|ID), family = "binomial" (link="logit"), data=joined_fooddata) | | | | |
|---|---|---|---|---|
| | Estimate | Std. Error | Z | P |
| Time | -0.33 | 0.68 | -0.49 | 0.62 |
| Treatment (exposed) | -1.66 | 0.79 | -2.11 | **0.035** |
| Sex (male) | -0.85 | 0.63 | -1.35 | 0.18 |
| Exposure group (two) | -0.30 | 0.56 | -0.53 | 0.60 |
| Time * Treatment | 1.55 | 1.01 | 1.52 | 0.13 |
| Posthoc contrasts: emmeans(model, pairwise ~ Treatment \| Condition) | | | | |
| | Estimate | Std. Error | Z | P |
| Before: Control vs Exposed | 1.66 | 0.79 | 2.11 | **0.035** |
| After: Control vs Exposed | 0.11 | 0.73 | 0.16 | 0.88 |

**Supplementary table S7**: Statistical model syntax and output for analysis of the proportion of time crayfish spent in the wastewater cue zone during the cue choice behavioural trial.

Effect of …. $N_{obs}$ = 688, $N_{crayfish}$ = 47

| glmmTMB(Proportion time in zone ~ Time bin + Treatment + Sex + (Experimental group * Cue side) + (Time bin \| ID), family="ordbeta", dispformula = ~ Treatment, data=joined_bindata) | | | | |
|---|---|---|---|---|
| | Estimate | Std. Error | Z | P |
| Time | 0.023 | 0.016 | 1.425 | 0.15 |
| Treatment (exposed) | -0.32 | 0.13 | -2.421 | **0.016** |
| Sex (male) | -0.20 | 0.15 | -1.357 | 0.17 |
| Experimental group (two) | 0.32 | 0.18 | 1.738 | 0.082 |
| Cue side (right) | 0.34 | 0.19 | 1.785 | 0.074 |
| Experimental group * Cue side | -0.65 | 0.26 | -2.521 | **0.012** |

**Supplementary table S8**: Statistical model syntax and output for analysis of activity rates across the three behaioural trials.

$N_{obs}$ = 141, $N_{crayfish}$ = 50.

| glmmTMB(Activity rate cm per min ~ (Treatment * Experiment) + Sex + Experimental group + Total length + (1\|ID), family = "gaussian", data = activitylong.na) | | | | |
|---|---|---|---|---|
| | Estimate | Std. Error | Z | P |
| Treatment (exposed) | 59.23 | 8.39 | 7.06 | **<0.0001** |
| Experiment (food) | 5.71 | 6.32 | 0.90 | 0.366479 |
| Experiment (shelter) | -40.17 | 6.32 | -6.35 | **<0.0001** |
| Sex (male) | -14.14 | 7. 22 | -1.96 | 0.050 |
| Experimental group (two) | -4.90 | 6.56 | -0.75 | 0.46 |
| Total length | 1.53 | 0.56 | 2.71 | 0.0067 |
| Treatment (exposed) * Experiment (Food) | -49.55 | 9.60 | 5.16 | **<0.0001** |
| Treatment (exposed) * Experiment (shelter) | -36.40 | 9.43 | 3.86 | **0.00011** |
| emmeans(model, pairwise ~ Treatment \| Experiment) | | | | |
| | Estimate | Std. Error | *t* | P |
| Choice: cont vs exposed | -59.24 | 8.39 | -7.061 | **<0.0001** |
| Food: cont vs. exposed | -9.69 | 8.56 | -1.133 | 0.26 |
| Shelter: cont vs. exposed | -22.83 | 8.37 | -2.73 | **0.0073** |

**Supplementary table S9**: Statistical model syntax and output for analysis of cholinesterase enzyme activity rates.

| glmmTMB(che ~ Treatment + Sex + Experimental group, family = "gaussian", data= joined_chedata) | | | | |
|---|---|---|---|---|
| | Estimate | Std. Error | Z | P |
| Treatment (exposed) | -0.17 | 0.079 | -2.13 | **0.032** |
| Sex (male) | 0.064 | 0.088 | 0.72 | 0.47 |
| Experimental group (two) | 0.046 | 0.081 | 0.57 | 0.57 |

**Supplementary table S10**: Statistical model syntax and output for analysis of the relationship between cholinesterase enzyme activity and crayfish activity rate across the three behavioural trial contexts.

| glmmTMB(Activity rate cm per minute ~ ChE activity + Total Length + (1\|ID), data=corr_data) | | | | |
|---|---|---|---|---|
| | Estimate | Std. Error | Z | P |
| ChE activity | -36.51 | 12.98 | -2.81 | **0.0049** |
| Total length | 0.59 | 0.67 | 0.88 | 0.38 |
